# Cancer-mediated Axonal Guidance of Sensory Neurons in a Microelectrode-based Innervation MPS

**DOI:** 10.1101/2023.10.18.562227

**Authors:** Matthijs van der Moolen, Andrea Lovera, Fulya Ersoy, Sacha Mommo, Peter Loskill, Paolo Cesare

## Abstract

Despite recent advances in the field of microphysiological systems (MPS), availability of models capable of mimicking the interactions between the nervous system and innervated tissues is still limited. This represents a significant challenge in identifying the underlying processes of various pathological conditions, including neuropathic, cardiovascular and metabolic disorders. In this study, we introduce a compartmentalized 3D coculture system that enables physiologically relevant tissue innervation while recording neuronal excitability. By integrating custom microelectrode arrays into tailored glass chips microfabricated via selective laser-etching, we developed an entirely novel class of innervation MPSs (INV-MPS). This INV-MPS allows for manipulation, visualization, and electrophysiological analysis of individual axons innervating complex 3D tissues. Here, we focused on sensory innervation of 3D tumor tissue as a model case study since cancer-induced pain represents a major unmet medical need. The system was compared with existing nociception models and successfully replicated axonal chemoattraction mediated by nerve growth factor (NGF). Remarkably, in the absence of NGF, 3D cancer spheroids cocultured in the adjacent compartment induced sensory neurons to consistently cross the separating barrier and establish fine innervation. Moreover, we observed that crossing sensory fibers could be chemically excited by distal application of known pain-inducing agonists only when cocultured with cancer cells. To our knowledge, this is the first system showcasing morphological and electrophysiological analysis of 3D-innervated tumor tissue *in vitro*, paving the way for a plethora of studies into innervation-related diseases and improving our understanding of underlying pathophysiology.

**Significance Statement:** MPSs integrating innervated tissues are crucial as alternatives to animal models and to enhance the translation of non-clinical research to humans, as nerve fibers play a pivotal role in many organs. To address this need, we developed the INV-MPS, enabling parallelized electrophysiological recording from 3D cocultures of peripheral neurons with complex tissues. We demonstrate the ability of cancer spheroids to direct axonal growth and modulate excitability of nociceptive fibers in a 3D setting. We anticipate that this model will be applicable for closely monitoring innervation in various tissues, including somatic and visceral organs. The INV-MPS, hence, has tremendous potential for human-relevant drug discovery, creating opportunities for more ethical and clinically translatable approaches to study innervation in disease.

## Introduction

MPS technology has made significant strides in enabling the investigation of complex biological interactions in accurately represented organs and tissues. However, the importance of innervation for proper tissue function has been largely overlooked [1]. Peripheral innervation plays an essential role in mediating the response of skin, bone, muscles and visceral organs to environmental changes and maintaining tissue homeostasis [2-4]. Additionally, in diseases such as cancer, aberrant innervation patterns can have significant effects on pain, highlighting the critical role of innervation in pathological conditions. Therefore, incorporating nerves in three-dimensional (3D) *in vitro* models is urgently required. Although previous approaches have implemented micropatterned structures to manipulate axons by injury and/or growth factor cocktails, these are limited to two-dimensional (2D) space and not appropriate for innervation of intricate 3D structures such as organoids and spheroids [5-10]. Additionally, these models lack electrophysiological readouts for monitoring the activity of 3D tissue’ innervating fibers and are mostly low throughput. To address these shortcomings, we developed a compartmentalized 9-well INV-MPS for coculture of peripheral nervous system (PNS) neurons with 3D tissues, all embedded in hydrogels. Additionally, integrated microelectrode arrays (MEA) allow for direct and parallelized recording in 3D neurons of electrophysiological activity, which is more physiologically relevant than calcium imaging and more efficient than patch-clamp recording [11]. As a proof-of-concept for the potential use of the INV-MPS for innervation studies, we specifically investigate sensory innervation of cancer spheroids. This is also motivated by the significant unmet medical needs associated with cancer-induced pain (CIP), as well as the extensive amount of literature available on nociceptive fibers, both *in vivo* and *in vitro,* and growing ethical concerns regarding *in vivo* animal research in this field.

The INV-MPS is fabricated by bonding a selectively laser-etched glass chip to a custom MEA, creating two individual compartments connected by micro-tunnels. Suspended in hydrogel, murine dorsal root ganglion (DRG) neurons and colorectal or pancreatic cancer cells are deposited into the respective compartments. Once axons extend through the barrier, interaction of neurites and cancer spheroids can be monitored via time-resolved live imaging and electrophysiological recording. Current literature, mainly based on *in vivo* studies, reveals an increase in nerve density/nerve alteration and pain states, also triggered by tumor-mediated release of factors such as NGF and prostaglandins (PGE2) [12-18]. Here we demonstrate cancer-mediated sensory fiber chemoattraction, thereby recapitulating these *in vivo* conditions within an *in vitro* setting. By exposing compartmentalized nerve terminals to pain and inflammatory mediators after applying a NGF gradient we confirmed the potent effect of NGF on inducing a nociceptive phenotype. Remarkably, in the absence of NGF, colorectal cancer spheroids were also able to attract sensory neurites and become innervated, accompanied by the induction of excitability through application of pain agonists.

These findings suggest that cancer cells alone can drive a nociceptive phenotype, similar to NGF. Finally, live-imaging of Ca^2+^ transients in cancer-innervating fibers confirmed that the neurites are responsive to noxious stimuli while interacting closely with cancer cells. Functional innervation of cancer spheroids by sensory fibers as shown here serves as a case study for the INV-MPS, an innovative 3D *in vitro* model amenable for exploring tissue-induced axonal growth, nerve sprouting and associated changes in neuronal electrophysiology. However, the platform has the potential to benefit a much broader community of microphysiological scientists who aim to recapitulate and explore the complexity of innervated tissues. Thus, the INV-MPS represents a significant advancement towards novel human-relevant drug discovery models, particularly in pathologies where tissue innervation plays a critical role.

## Results

### A novel MPS with integrated electrodes for 3D axon manipulation and recording from sensory neuron cultures

Although MPSs have advanced with rapid strides recently, the development of new in vitro PNS models has been hampered by challenges in integrating effective readouts into complex 3D cultures. However, improving existing models, particularly with respect to tissue innervation, is critical for studying pathophysiological processes and the development of new therapeutic strategies. Therefore, we developed a novel MPS that integrates a non-invasive electrophysiological readout with glass microfluidics to produce a valuable 3D neurite outgrowth model that separates soma and axons. Specifically, we grew peripheral neurons in a 3D scaffold and facilitated their projection through a physical barrier, enabling them to innervate axonally residing tissue (Fig. 1A and Movie S1). To achieve this, we produced a fused silica MPS patterned with µm-scale features and attached it to a microelectrode array substrate, as previously described [11]. The MPS includes pairs of adjacent two-layer tissue wells (somatic and axonal compartment) featuring defined hydrogel layers (GL; approx. 8.0 µL) below media layers (ML; approx. 110.0 µL). The two adjacent tissue wells are separated by a 300.0 µm thin barrier featuring 12 rows of 110 microtunnels (10.0 x 10.0 µm cross-section) each, over a surface area of approx. 4.0 x 1.0 mm (Fig. 1B and SI Appendix, Fig. S1A). This 3D-arrangement of µ-tunnels is possible due to the unique capabilities of selective laser etching (SLE) that enables the selective removal of glass from a monolithic piece in 3D. The microtunnels (Fig. 1C) allow only DRG neurites to pass into the GL of the axonal compartment and constrain DRG cell bodies to the somatic compartment. To record from gel-embedded 3D neurons inside the INV-MPS, we utilized capped microelectrodes (CMEs) positioned at the base of the somatic compartment that effectively trap axons (Fig. 1D). Neurons occupying CME-microtunnels also extended towards the barrier and crossed into the axonal compartment in 3D.

**Fig. 1.**
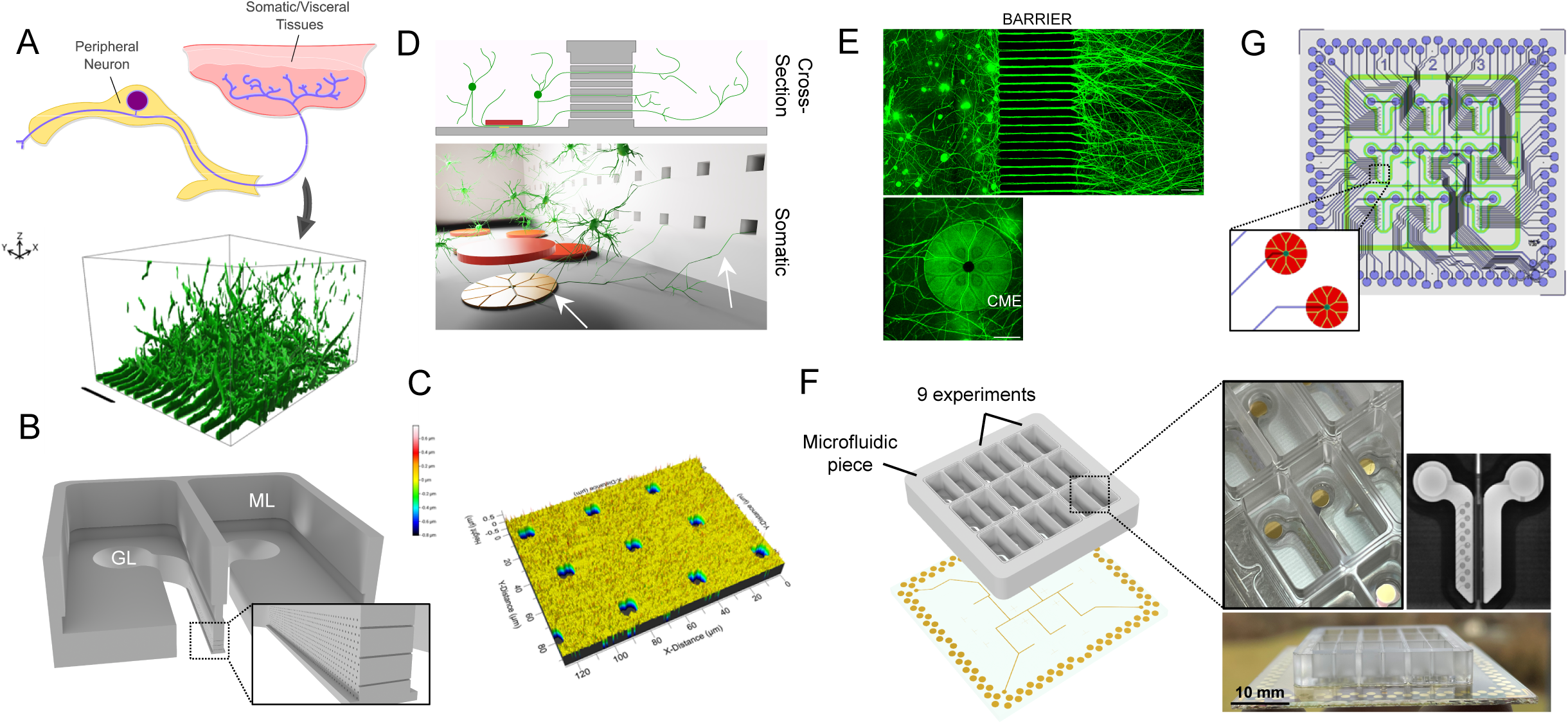
A novel MPS with integrated electrodes for 3D axon manipulation and recording from sensory neuron cultures. (A) Recapitulating the innervation of soft tissues by the peripheral nervous system. 3D-surface rendering of DRG terminals extending into 3D space in the axonal compartment. Dimensions 600 × 450 × 300 µm. (Scale bar: 100 μm.). (B) Both somatic and axonal compartments are filled with a soft 3D scaffold inside the gel layer (GL). Supporting media is added on top in both media layers (ML) (width: 300.0 μm.). (C) High magnification color map of micro tunneled barrier profilometry data. (D) DRG neurons grow neurites in 3D through the barrier while simultaneously projecting neurites into recording CMEs residing in the somatic compartment. (E) Live-cell confocal images (max projection) of GFP-transduced DRG homogeneously distributed across the somatic compartment with neurites traversing the barrier and entering CMEs (Scale bar: 75 μm.). (F) INV-MPS holds 9 identical cocultures with 13 CMEs each and assembles by bonding on top of the dedicated 120-electrode MEA. (G) Technical drawings of MEA-CME production photomasks.

To confirm the spatial separation of soma and nerve terminals, we used live-cell confocal fluorescence microscopy of DRG neurons transduced with adenoviral vectors encoding enhanced green fluorescent protein (eGFP) (Fig. 1E and Movies S2 and S3). Additionally, we confirmed that neurites grew inside the confined spaces of the CMEs (5.0 x 3.0 µm), allowing us to detect electrophysiological signals (up to ∼100 µV) generated upon firing of action potentials (SI Appendix, Fig. S1B). Each micropatterned glass chip was bonded onto a 49.0 x 49.0 mm glass substrate containing 117 embedded recording electrodes. Thirteen recording-CMEs and a reference electrode were positioned within each of the nine somatic compartments. CMEs were connected to edge-pads, which secured the connection to pins of the MEA-recording system (Fig. 1F and G).

### 3D axonal guidance and nerve terminal excitability modulated by NGF inside the MPS

Our study examined the responsiveness of 3D gel-embedded DRG neurons to neurotrophic gradients, which may have implications for tissue innervation under various pathophysiological conditions *in vivo*. We focused on NGF, a well-known neurotrophic factor that promotes the development of neuronal growth cones and can enhance neuronal survival *in vitro* [19, 20]. Here, experiments were performed using an INV-MPS without integrated electrodes (SLE INV-MPS; SI Appendix, Fig. S2C), which features a single row of tunnels at the bottom of the barrier, allowing for accurate comparison of neurite outgrowth and microtunnel diffusion kinetics. To analyze the impact of NGF on 3D sensory fiber’ extension into the axonal compartment, we supplemented NGF solely axonally (NGF -/+), thereby creating a stable gradient over the barrier for 1 week. We found a significant increase in fibers crossing the barrier (Fig. 2A and B and SI Appendix, Fig. S2A) with limited variability in survival across different conditions (SI Appendix, Fig. S2B). When NGF was supplied to both compartments (NGF +/+), few axons extended through the barrier and neurites predominantly grew within the somatic compartment, likely due to the lack of a chemoattracting gradient. To mimic a NGF gradient across adjacent wells, we monitored the concentration of fluorescently labelled dextran moieties (applied to the axonal compartment) for 12 hours and observed a ∼10x lower concentration of both 4 kDa and 20 kDa tracers in the somatic compartment. It is important to note that the small amount of dye that diffuses through the barrier is progressively diluted into a much larger volume in the somatic compartment, resulting in a significantly lower bulk concentration. Having optimized the conditions for neurite outgrowth in the SLE INV-MPS, we then transitioned to using the INV-MPS with integrated MEA for electrophysiological characterization of 3D sensory terminals in the NGF -/+ condition (Fig. 2D). Application of TRPV-1 agonist capsaicin [21] to the axonal compartment of the INV-MPS resulted in measurable action potentials in the majority of CMEs located in the somatic compartment (Fig. 2E). This provides evidence that compartmentalized 3D neuronal culture in the INV-MPS can successfully replicate peripheral activation of both receptors and ion channels involved in generating and transmitting nociceptive signals from nerve terminals to the soma.

**Fig. 2.**
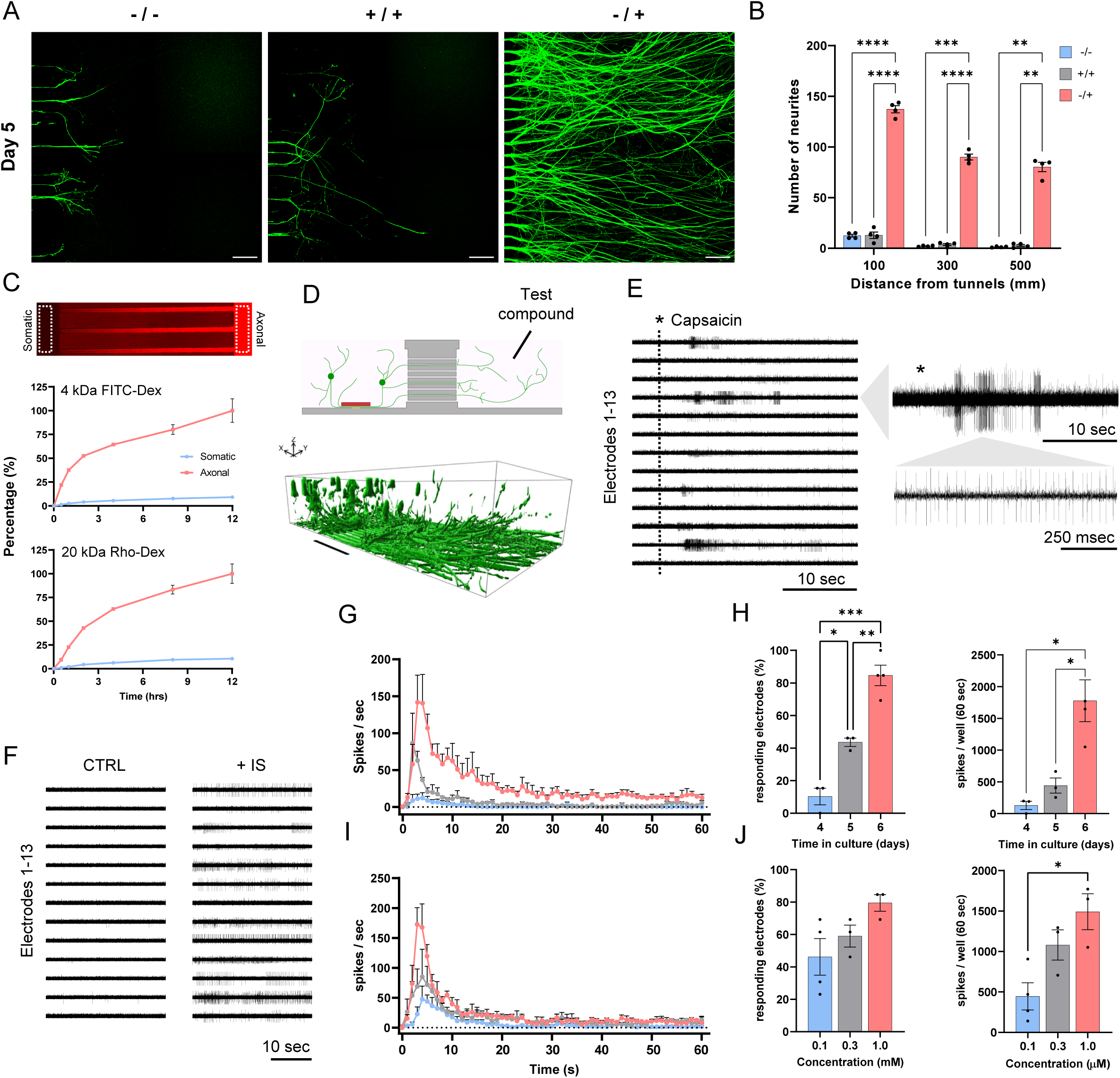
3D axonal guidance and nerve terminal excitability modulated by NGF inside the MPS. (A) Live-cell confocal images of GFP-transduced DRG neurons protruding from the barrier in 3D after 5 days in culture. Neurobasal-A culture medium either contains no NGF (-/-), NGF in both somatic and axonal media compartment (+/+) or NGF only in the axonal media compartment (-/+) (Scale bar: 75 μm.) (B) Quantification of neurite outgrowth after 5 days in culture. Total number of neurites was measured at 100 µm, 300 µm, and 500 µm from the barrier. Bar graphs represent mean ± SEM (n=4 wells) (C) Micrograph shows diffusion of Dextran-FITC (MW: 4 kDa) or Dextran-Rhodamine B (MW: 20 kDa) through the micro-tunnel barrier from axonal (right) to somatic (left). Fluorescent intensity profiles generated from ROIs taken 5 µm from the micro-tunnel barrier, plotted against time as percentages (normalized to intensity at T = 12 hrs). Data is represented as mean ± SEM (n=3 wells). (D) Agonists are applied to the media layer atop the 3D nerve terminals of a NGF -/+ culture in the axonal compartment. 3D-surface rendering of DRG neurons and their soma. Dimensions 1450 × 750 × 300 µm. (Scale bar: 300 μm.) (E) Trace plots representing the activity recorded by 13 CMEs after application of 1 µM Capsaicin (black asterisk) at different time scales. (F) Trace plots comparing control to activity recorded 5 minutes after application of inflammatory soup (IS containing: 3.0 µM Bradykinin, 3.0. µM Prostaglandin, 3.0. µM Serotonin and 3.0 µM Histamine). (G) Temporal dynamics of Capsaicin (1 µM) responses in -/+ cultures of DRG neurons after 4 (blue), 5 (grey) and 6 (red) days. Dots represent the mean spikes per second per well over a 60-second span post-application (n=3-4 wells). (H) Bar graphs comparing the development of nociceptive phenotype (percentage of responding electrodes) and total spikes recorded per well in the 60-second span post application at different time points (time in culture). (I) Dose-response plots comparing temporal dynamics following application of 0.1 (blue), 0.3 (grey) and 1.0 (red) µM capsaicin in -/+ cultures of DRG neurons (n=3-4 wells). (J) Bar graphs comparing CMEs excitability probability (percentage of responding electrodes) and total spikes recorded per well in the 60-second span post application following application of different capsaicińs concentrations. In B, H, and J, significant differences between mean values of groups are displayed as asterisks: ∗p < 0.05; ∗∗p < 0.01; ∗∗∗p < 0.001; ∗∗∗∗p < 0.0001; Brown-Forsythe and Welch’s ANOVA test.

Recorded signals appeared within a timeframe of 3-5 seconds from administration of capsaicin, indicating the system’s ability to rapidly detect and measure terminally-evoked changes in neuronal activity. Given the non-invasive nature of MEA electrophysiological recordings, our system enables investigation into the long-term effects of substances within an inflammatory milieu. This was demonstrated by recording changes in neuronal activity induced by an inflammatory mediator cocktail (bradykinin, prostaglandin, serotonin and histamine), where action potentials could be continuously measured for up to 30 minutes by the majority of electrodes (Fig. 2F). Our findings indicate that the INV-MPS can continuously monitor activity generated in 3D nerve terminals, providing a valuable tool for studying the mechanisms involved in peripheral inflammatory pain. Examining electrode response rates following capsaicin application, we found that a response rate of over 80% can be achieved within 6 days from plating. This is longer compared to typical MEA recordings of DRG cultures in 2D (2-3 days), most likely due to the additional time required for neurites to extend into the axonal compartment (Fig. 2G and H). Application of different doses of capsaicin (0.1-1.0 µM) on day 6 revealed a direct relationship between the concentration of the agonist and the number of observed responses and their potency (Fig. 2I and J), suggesting that the INV-MPS could facilitate applications in drug discovery. In summary, the INV-MPS could be utilized to study the engagement of specific ligand-gated receptors or voltage-gated ion channels that may play a role in modulating pain states, as well as to monitor the potential of various factors to induce neuronal outgrowth or nerve degeneration in toxicity studies.

### Induction of sensory neurite outgrowth by colorectal cancer spheroids in the absence of NGF supplementation

We then examined whether the INV-MPS could be utilized as a tool for evaluating changes in nerve density and innervation induced by cancer in a neuron-cancer coculture. This investigation could contribute to the comprehension of the underlying neuronal mechanisms involved in pathological pain *in vivo* [22-25]. We seeded DRG neurons in the somatic compartments of the SLE INV-MPS, while colorectal (HT29) or pancreatic (PANC1) cancer spheroids were seeded in the axonal compartment. We monitored neurites as they traversed the barrier and innervated axonal targets following application of NGF to the distal compartment (NGF -/+). No cancer-mediated effects on neuronal outgrowth were observed compared to the NGF -/+ control condition (Fig. 3A and B). This suggests that NGF in the media of the axonal compartment may saturate the outgrowth potential of nerve fibers and override guidance clues. Therefore, we opted to omit NGF from the cocultures (NGF -/-) and were then able to demonstrate a cancer-mediated increase in nerve fiber density in the axonal compartment of HT29 - DRG cocultures (Fig. 3C and D). Remarkably, we were able to simultaneously demonstrate innervation and complete envelopment of individual cancer spheroids, as observed *in vivo* [13, 16].

**Fig. 3.**
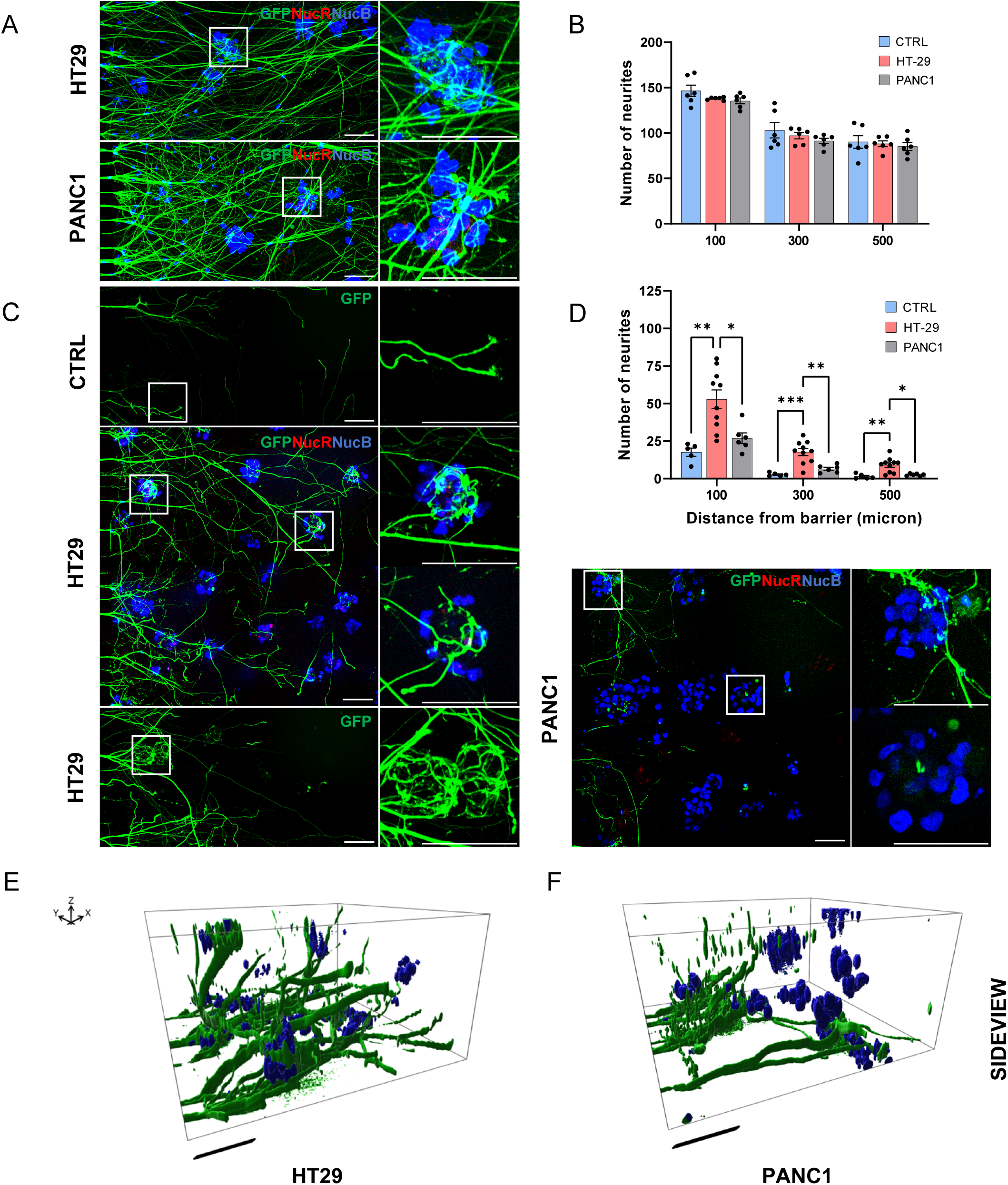
Induction of sensory neurite outgrowth by colorectal cancer spheroids in the absence of NGF supplementation. (A) Live-cell confocal images of GFP-transduced DRG neurons protruding from the micro tunnel-barrier and innervating either HT29 (colorectal) or PANC1 (pancreatic) cancer spheroids in 3D after 6 days in a NGF -/+ culture. Cancer spheroids are visualized using dyes staining nuclei (NucB) and dead cells (NucR). (Scale bar: 75 μm.) (B) Quantification of neurite outgrowth after 6 days in culture. CTRL represents a DRG-only (NGF -/+) culture at day 5 & 6. Bar graphs represent mean ± SEM (n=6 wells) (C) Live-cell confocal images of DRG neurons protruding from the micro tunnel-barrier and innervating either HT29 or PANC1 cancer spheroids after 6 days in a NGF -/- culture. (D) Quantification of neurite outgrowth after 6 days in culture. CTRL represents a DRG-only (NGF -/-) culture (n=5-10 wells). (E) Sideview of 3D-surface renderings of sensory neurites innervating HT29 or (F) PANC1 cancer spheroids. Dimensions 400 x 400 x 200 µm. (Scale bar: 100 μm.) In B and D, significant differences between mean values of groups are displayed as asterisks: ∗p < 0.05; ∗∗p < 0.01; ∗∗∗p < 0.001; ∗∗∗∗p < 0.0001; Brown-Forsythe and Welch’s ANOVA test.

3D representations of nerve fibers extending into the axonal compartments among resident cancer spheroids further confirm that the described cell-cell interactions occur in all dimensions (Fig. 3E and F and Movies S4, S5, S6 and S7). Furthermore, for optimal comparison of neuronal outgrowth conditions and standardization, cancer cells were cultivated in neuronal media instead of cancer media without clear negative effects on viability of the cancer spheroids (SI Appendix, Fig. S3A and B). Interestingly, while HT29 - DRG cocultures exhibited a significant enhancement of neuronal outgrowth compared to control conditions (DRG alone, NGF -/-), this effect was not observed in PANC1 - DRG cocultures (Fig. 3C and D). This observation suggests that innervation can be highly cell-type-specific, possibly due to the presence of different molecular or paracrine cues associated to the microenvironment of each cancer type.

### Sensory neurons innervating 3D colorectal cancer spheroids can be activated by pain mediators

The combination of electrical and image-based techniques is crucial for investigating neuronal functions. However, current methods for monitoring activity in 3D tissues, such as calcium imaging, are typically limited to qualitative analysis. In this study, we demonstrate the potential of the MEA-integrated INV-MPS for screening purposes in pain research. As shown in Fig. 3, cancer spheroids occupying the axonal compartment of the SLE INV-MPS can stimulate nerve growth, raising the question of whether nerve terminals also exhibit excitability (Movie S8). To address this, we applied either capsaicin or bradykinin to the axonal compartment of INV-MPSs containing DRG (NGF-/-) with or without cancer spheroids. Prior to chemical excitation, we confirmed that HT29 did not cause any rise in spontaneous activity (SI Appendix, Fig. S4A). In the presence of cancer spheroids it was possible to measure characteristic responses to both compounds in axonally extended nerve fibers, which were absent in the control group (Fig. 4A and B). Although the responses were less pronounced than those evoked in DRG monocultures (NGF-/+; Fig. 2E-J), our data suggests that the inclusion of HT29 cancer spheroids not only drives neurite outgrowth, but it is sufficient to induce nerve terminal excitation in response to chemical stimuli. The response to both depolarizing stimuli was quantified and compared between DRG and HT29 coculture vs. DRG control (Fig. 4C & D and SI Appendix, Fig. S4B). The results show that capsaicin application to HT29 coculture induced responses characterized by a less pronounced initial cumulative peak and lower responsiveness compared to NGF-/+ conditions (Fig. 2I and J), which is consistent with the difference in neurite outgrowth (<50%). Deeper analysis, including time-course and number of responses elicited by capsaicin or bradykinin applied to DRG (NGF-/-) alone or cocultured with HT29 cancer spheroids, revealed that colorectal cells have the capacity to enhance both outgrowth of 3D nerve terminals and transduction of noxious stimuli into action potentials.

**Fig. 4.**
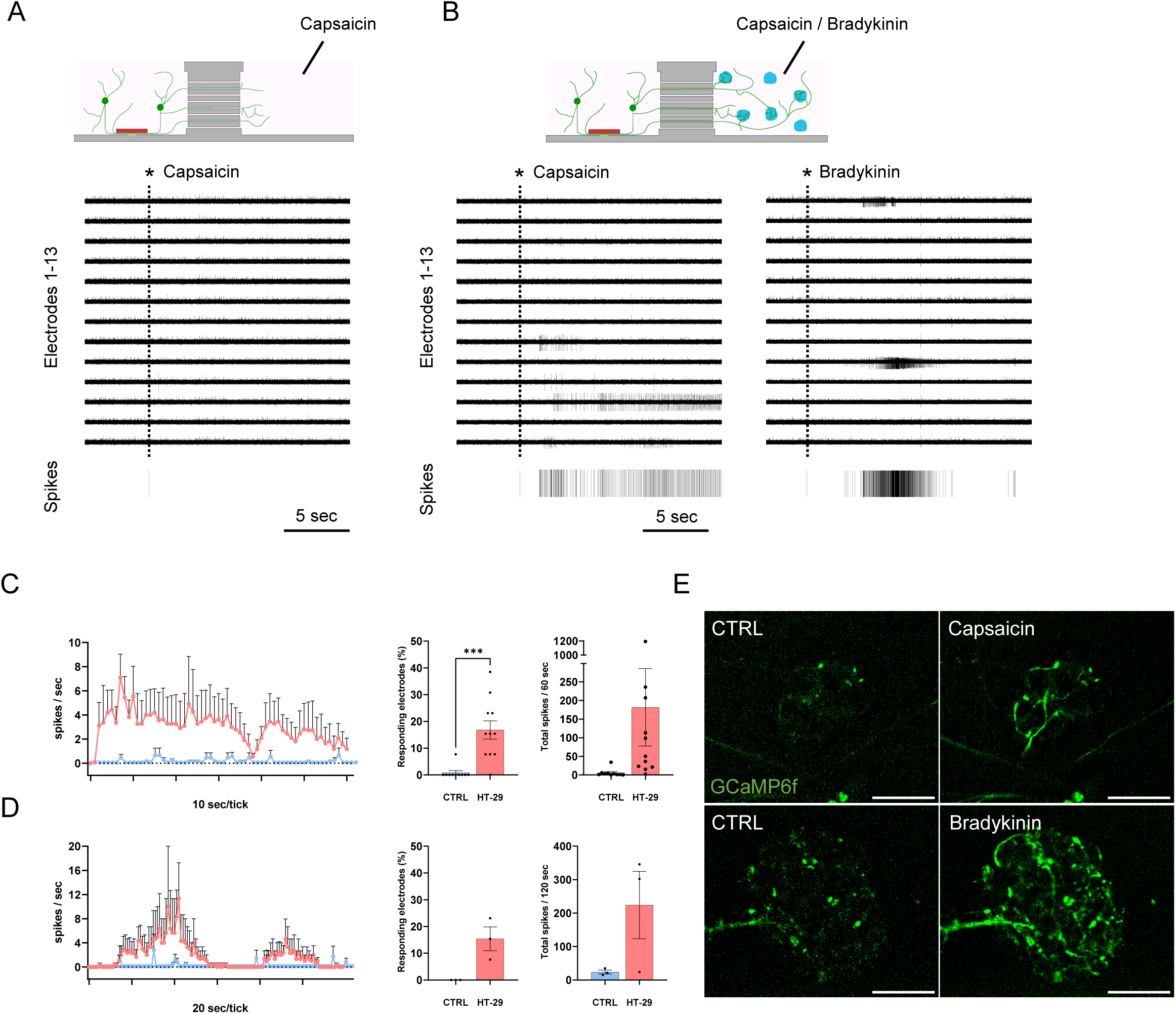
Sensory neurons innervating 3D colorectal cancer spheroids can be activated by pain mediators. (A) Trace plots representing individual CMEs (1 to 13) and cumulative raster after application of 1.0 µM Capsaicin (black asterisk) to a DRG-only culture (NGF -/-) after 6 days. (B) Trace plots and cumulative raster after application of 1.0 µM Capsaicin or 1.0 µM Bradykinin to DRG neurons innervating HT29 cancer spheroids (NGF -/-) after 6 days. (C) Temporal dynamics of capsaicin responses in DRG neurons innervating HT29 cancer spheroids (NGF -/-). Dots represent the mean spikes per second per well over a 60-second span post-application (n =10-11 wells). Bar graphs comparing excitability of DRG neurons responding to axonal stimuli (percentage of responding electrodes) and total spikes recorded per well in the 60-second span post application. (D) Temporal dynamics of bradykinin responses in NGF -/- cultures of DRG neurons innervating HT29 cancer spheroids. Dots represent the mean spikes per second per well over a 120-second span post-application (n=3 wells). Bar graphs comparing excitability of DRG neurons responding to axonal stimuli (percentage of responding electrodes) and total spikes recorded per well in the 120-second span post application. (E) Live-cell confocal images of GCaMP6f-transduced DRG neurons innervating HT29 cancer spheroids after 6 days in a NGF -/- culture. Application of 1.0 µM Capsaicin and 4.0 µM Bradykinin induces Ca^2+^ transients (also seen in Movie S9 & S10). In C and D, significant differences between mean values of groups are displayed as asterisks: ∗p < 0.05; ∗∗p < 0.01; ∗∗∗p < 0.001; ∗∗∗∗p < 0.0001; Brown-Forsythe and Welch’s ANOVA test.

To explore a more direct association between nerve terminal excitability and cancer innervation, we employed AAV-GCaMPf6 to directly visualize calcium transients induced by application of chemical agonists in DRG neurons innervating cancer spheroids (Fig. 4E and Movies S9 and S10). Although Ca^2+^ imaging can indeed provide an optical measure of nerve terminal activation, monitoring responses in a multiplexed 3D setting remains technically challenging and time-consuming, which emphasizes the advantages of the INV-MPS platform for quantifying electrical excitability in 3D microtissues.

## Discussion

The MPS-field aims to replicate human organ functions not properly mimicked by conventional cell cultures or animal models [26]. However, MPSs for the study of organ innervation are currently not available due to the added complexity of incorporating nerve fibers, despite the potential benefits for therapeutic domains such as visceral pain, metabolic and cardiovascular disease and neurological disorders [27]. Novel designs and platforms are required to integrate peripheral neurons and target tissues, whilst providing a robust electrophysiological readout that is consistent with neuronal activity. By accurately monitoring and controlling electrical activity of innervating fibers, these will aid the understanding of underlying mechanisms that drive peripheral neuron-tissue interaction leading to pathological phenotypes. Leveraging expertise in combining custom MEA, high-resolution laser glass etching and 3D neuronal cultures [11], we developed an innervation platform that meets the requirements of the MPS-field towards creating intricate tissue models and incorporates a reliable electrophysiological readout. Unlike conventional 2D axonal outgrowth models [28], the INV-MPS is used to closely mimic 3D *in vivo*-like nerve-tissue interplay and simultaneously overcomes the limitations of conventional MEA systems solely designed for recording from 2D cultures. This is achieved by integrating capped microelectrodes that allow electrophysiological recording from 3D peripheral nerve fibers innervating hydrogel-embedded target tissues. Distinct from alternative methods such as patch-clamp, our approach presents several unique features, including non-invasive readout, enhanced throughput, simple handling, and seamless integration across platforms. Notably, the INV-MPS avoids the utilization of patterned polymers such as polydimethylsiloxane (PDMS), a common choice for microfluidic devices. This is due to PDMS’s susceptibility to small molecule absorption and the intricate molding procedures it entails, among other factors. Instead, we designed an easy-to-assemble, reusable device utilizing a glass microfluidic chip and MEA to compartmentalize and record from cultured sensory neurons, effectively ensuring the required separation of soma and terminals through barrier-etched microtunnels [5]. Axonal growth through the barrier was shown to strongly depend on a NGF gradient and uniformly penetrates the 3D hydrogel-substrate, closely recapitulating physiologically relevant nerve guidance [29].

Importantly, we demonstrated that our platform enables non-invasive measurement of the electrical activity of 3D sensory terminals and that the response profiles to pain mediators are similar to those observed *in vivo* and *ex vivo* [30, 31]. These results suggest that the INV-MPS can provide a physiological relevant measure of the engagement of receptors and ion channels localized at distinct locations (terminal vs. somatic) potentially mediating nociception. Simultaneously, it allows for the monitoring of neurotrophic factors/inhibitors’ effects on neuronal outgrowth and arborization, providing valuable insights into these processes. Next, we aimed to apply INV-MPS’ unique readout capabilities to the study of disease-related phenotypes by exploring CIP. Our findings indicate that when cancer spheroids and DRG neurons are cocultured in the presence of NGF, neural outgrowth is primarily driven by the exogenously applied neurotrophin pool. Remarkably, we observed that when NGF is omitted, the presence of colorectal cancer spheroids is sufficient to stimulate neural outgrowth and ultimately the innervation of individual cancer structures, as observed *in vivo* [13, 16]. This emphasizes the complex interplay between cancer and the PNS, highlighting the need to consider the impact of exogenously applied NGF or other growth factors when studying nerves and nerve-tissue crosstalk *in vitro*. Understanding the role of released factors is crucial to elucidate underlying mechanisms of chronic pain and identify potential therapeutic targets for the management of CIP. An increase in nerve fiber density could be demonstrated for colorectal cancer spheroids (HT29) but not for pancreatic cancer spheroids (PANC1). However, it would be premature to conclude that pancreatic cancer cannot induce outgrowth within the INV-MPS setting, as induction of axonal growth can have varying degrees of dependency on tumor-associated stromal cells and inflammation [13, 32-36], only partially recapitulated here. Neuronal outgrowth was clearly shown to depend on exogenously applied NGF, suggesting a potential correlation between the observed cancer-mediated outgrowth and NGF presence. However, given that both HT29 and PANC1 release comparable amounts of NGF *in vitro* [37, 38], the observed disparity in innervation between these two cancer types poses that NGF release alone may not be sufficient to drive outgrowth in this specific *in vitro* context. Expanding on this by investigating putative paracrine mediators and exploring alterations in gene expression and regulation of relevant protein levels is beyond the scope of our current study. However, the observed difference in 3D innervation between colorectal and pancreatic cancer spheroids emphasizes the ability of the INV-MPS to accurately recapitulate and detect differences in cancer-specific environments. This was further assessed by measuring the electrophysiological response to prototypic noxious and inflammatory stimuli in innervating fibers of HT29 - DRG neuron cocultures. Here, nerve terminals extending into the axonal compartment and enveloping cancer spheroids were found to respond to both capsaicin and bradykinin. Measured responses were remarkably greater (duration and frequency) than those observed in cultures without cancer and exogenously applied NGF.

This was complemented by live-imaging of evoked calcium transients occurring in 3D cancer-innervating fibers, providing a direct qualitative evaluation of tumor innervation and neuronal activity. Having the ability to additionally image nerve terminals and cancer cells in direct contact suggests that the INV-MPS holds potential as a valuable multimodal platform for investigating the reciprocal influences between cancer and closely associated innervating fibers. While the primary objective of this study was to validate the INV-MPS by examining the impact of cancer microtissues on sensory innervation and excitability, emerging evidence indicates that peripheral neurons may additionally influence cancer homeostasis and dissemination [39], making it a promising avenue for future exploration using this *in vitro* model. The potential applications of the INV-MPS extend beyond CIP and cancer, as this model offers the opportunity to investigate a diverse array of diseases by faithfully reproducing 3D peripheral innervation and measuring excitability in various tissues. In conclusion, the INV-MPS is purposefully designed to facilitate the exploration of complex *in vitro* biology, presenting great potential for researchers in the MPS field who seek to understand the intricate interactions between the PNS and innervated organs.

## Materials and Methods

### Animals

Wild-type Swiss mice obtained from Janvier Labs (France) were housed in the animal facility of the institute. Mice were kept under standard conditions with controlled temperature and a 12-hour light-dark cycle, with ad libitum access to food and water. All experimental procedures described below were performed in compliance with the regulations outlined in the European Union (EU) legislation for the care and use of laboratory animals (Directive 2010/63/EU of the European Parliament on the protection of animals used for scientific purposes) and the German TierSchG (Tierschutzgesetz) with the latest revision in 2019.

### Microfabrication

INV-MPS devices were prepared following the protocol described previously [14], with modifications. Briefly, compartmentalized microfluidic chips (see SI Appendix, Fig. S1A) were fabricated by SLE using fused silica. MEAs with integrated CME were produced by structuring two layers of permanent epoxy photoresist on gold microelectrodes. SU-8 2002 (MicroChem, USA) was spin-coated to achieve a thickness of approximately 3.0 µm (10 s at 500 rpm, then 30 s at 1000 rpm) (see SI Appendix, Fig. S1B). ADEX TDFS A20 (Micro Resist Technology GmbH, Germany) was then laminated onto the SU-8 layer using a pouch laminator (GMP Photonex@325, EF02015) at a temperature of 75°C. Microfluidic chips were bonded to the MEA using EPO-TEK 301-2FL (Epoxy Technology) after alignment with a Fineplacer® lambda (Finetech GmbH, Germany). The bonded devices were cured at 80°C for 3 hours. Prior to use, the devices were rinsed with bi-distilled water and incubated in a 1% Tergazyme solution for 3 hours. Subsequently, they were washed overnight with bi-distilled water. This cleaning process was repeated between experiments if the devices were deemed suitable for reuse.

Alternatively, the devices were detached from the MEA substrate by immersion in concentrated sulfuric acid (H2SO4 ROTIPURAN® 98% - Carl Roth, # H290-H314) overnight and then reattached to a new MEA substrate. SLE INV-MPS devices do not acquire additional assembly as they are created from a monolithic fused silica substrate and lack microelectrodes.

### Fused silica microfluidic chip production

Fabrication of fused silica microfluidic chips is based on a subtractive, direct-write microfabrication process. Exploiting a femtosecond laser and etching creates integrated 3D micro-systems with micrometer precision [40]. The process exploits highly localized material modifications, triggered by non-linear absorption during the laser exposure step. The size of the modified region is in the order of 2.0 µm in XY-axis and from 10.0 µm to 100.0 µm in Z-axis, depending on the optics used to focus the laser. These modifications cause a local increase of ‘etching rate’ in the exposed material. Since modification happens only within the focal spot (voxel) e.g., buried structures or free-form surfaces can be produced. The exposed material is later etched in KOH based solution to create 3D devices.

### Profilometry

Barrier integrated microtunnels were analyzed using a 3D optical profiler (Contour GTK-A, Bruker UK Ltd.) equipped white light illumination and a 10x magnification objective. The instrument provides lateral resolution of 1.0 µm and Z resolution below 10.0 nm. Stylus profilometry (DektakXT, Bruker UK Ltd.) was performed to characterize CME height. Both SU8 (no ADEX cap) and SU8-ADEX (with cap) were measured by using a 2.0 µm tip. Both sets of measurements were processed and analyzed using dedicated Vision64 software (Bruker UK Ltd.).

### 3D DRG neuron culture

Dorsal root ganglion neurons were isolated and dissociated by modification of a previously published protocol [41]. Postnatal mice (P4-6) were sacrificed, in accordance with the European Union (EU) legislation, by decapitation with dissection scissors and disinfected with ethanol. Narrow incision through the skin and muscle of the back then exposed the spine, after which the ganglia could be removed using forceps for preservation in Hibernate™-A Medium (Thermo Fisher, #A1247501). Cleaned ganglia were then incubated in enzymatic solution(s) containing collagenase (IV) (Worthington, #LS004186) and dispase (Worthington, #LS02109), followed by manual dissociation in DNAse solution (Worthington, #LS002139). Lastly, gradient centrifugation was performed to remove dead cells and debris before mixing with plating media Neurobasal-A (Thermo Fisher, #10888-022) with GlutaMAX (Thermo Fisher, #35050-038), SM1 (Stemcell, #5711), N2 Supplement-A (Stemcell, #7152), 0.5% Penicillin-Streptomycin (Sigma, #P4333) and w/wo 100 ng/mL Nerve Growth Factor 2.5S (Promega, #G5141).

Cell suspension was counted using a NucleoCounter NC-200 (ChemoMetec A/S, Denmark) and mixed with Matrigel® Growth Factor Reduced (Corning, #356230) at 1:4 ratio, resulting in a complex modulus of 20 ± 2 Pa (mean ± SD), and kept in a cold rack to prevent matrigel polymerization during seeding. Up to 5 INV-MPS devices were simultaneously sterilized / hydrophilized by air plasma treatment (Piccolo, Plasma Electronic GmbH). Low retention tips were used to dispense 8.5 µL of cell solution into each proximal channel of a single compartmentalized experiment (gel layer 1000.0 µm thick - 9 total), followed by similar filling of the distal channel w/wo cancer cells. DRG neurons were plated at RT with stand. conc. 800 cells/µL (3200 cells/µL before mixing) and left to polymerize before addition of 100 µL Neurobasal-A to the media layer. 3D cultures were maintained up to 1 week by adding fresh Neurobasal-A medium every day (50/50) until electrophysiological or imaging readout.

### AAV1-eGFP and AAV1-GCaMP6f transduction

DRG neurons cultured in 3D were transduced immediately after plating with adeno-associated virus serotype 1 (AAV1) vectors containing either a transgene encoding enhanced green fluorescent protein (eGFP) or a G-protein coupled calcium indicator (GCaMP6f), both under control of the human Synapsin 1 (hSyn) promoter. Viral particles pAAV-hSynEGFP and pAAV-hSyn-GCaMP6f.WPRE.SV40, were obtained from Addgene (catalog #50465 and #100837, respectively). The multiplicity of infection (MOI) used was 105. After 3-4 days in culture, DRG neurons exhibited homogeneous green fluorescence in the case of eGFP, while GCaMP6f-labeled DRG neurons showed fluorescence specifically upon Ca^2+^ transients.

### Cancer spheroid culture preparation

Human colorectal adenocarcinoma (ATCC, #SK809) and human pancreatic carcinoma (ATCC, #CRL1469) cells were grown in McCoy’s 5A medium (Gibco, #16600082) and Dulbecco’s Modified Eagle’s medium (Gibco, #11965092), respectively, both supplemented with 10% Fetal Bovine Serum (Gibco, 10499044) and 1% Penicillin-Streptomycin (Sigma, #P4333). Cells were seeded onto uncoated T75 flasks at a density of 1.0 × 106 cells per flask. On plating day, cells were detached with Trypsin-EDTA 0.25% (Sigma, T4049) for 3 min at 37 ◦C, enzymatic digestion was then blocked using DMEM (Gibco, 31966-021) with 10% FBS and cells were spun down before resuspending in complementary cancer media or Neurobasal-A for plating. As described in “3D DRG neuron culture”, cell suspensions were similarly counted using a NucleoCounter NC-200, mixed with Matrigel 1:4 for stand. conc. of 500 cells/µL (2000 cells/µL before mixing) and kept cold while preparing INV-MPS devices. For neuron-cancer coculture experiments – after dispensing 8.5 µL of DRG solution into the proximal compartment, either HT29 or PANC1 cell solution was dispensed into the distal compartment at RT and left to polymerize to ensure optimal 3D distribution.

100 µL Neurobasal-A w/wo NGF (100 ng/mL) (or cancer media for live-dead assays) was then added on top in the media layer and cells were closely monitored throughout the week to qualify their aggregation into cancer spheroids. 3D cultures were maintained up to 1 week, by adding fresh medium every day (50/50) until electrophysiological or imaging readout.

### Fluorescence imaging of neurite growth and innervation

3D structures were imaged by a spinning disk microscope (Zeiss Cell Observer®) equipped with ZEN software (blue edition, ZEISS), followed by quantitative analysis using open-source image processing software, FIJI (ImageJ, http://imagej.nih.gov/ij/) or qualitative reconstruction with IMARIS 4.6 (Bitplane AG, Oxford Instruments). A dedicated 384 well microplate adaptor was designed and produced by Weerg Srl (Italy) to simultaneously hold two devices while imaging under humidity/CO2/temperature control (SI Appendix, Fig.S2B). Neurite outgrowth assays (SI Appendix, Fig.S2C) were performed on maximum intensity projections of 100 slices (AAV1-eGFP Alexa Fluor 488 - z-axis with 0,75 µm spacing – 20x) for 3 randomly selected ROIs (700 µm x 700 µm) starting at substrate-height. For each ROI, intensity profiles of vertical lines were plotted at 100 µm, 300 µm and 500 µm distance from distal tunnel exits and a ‘Find Peaks’ algorithm was then applied that returns the number of maxima (representing neurites – after background subtraction) (available from https://github.com/tferr/Scripts#ij-bar/). Neuron-cancer cocultures were quantified for neuronal outgrowth by identical image generation and processing with additional pre-incubation (3 drops/mL for 90 min at 37◦C) of NucRed Dead 647 ReadyProbes™ Reagent (TO-PRO-3 iodide) (Invitrogen, #R37113) for dead cells and NucBlue Live ReadyProbes™ Reagent (Hoechst 33342) (Thermo Fisher, #R37605) for all nuclei.

### Dextran diffusion

Barrier permeability assay was performed using 2 different fluorescently labelled dextran moieties. 4 kDa FITC-Dextran and 20 kDa Rhodamine-Dextran (100 µg/mL diluted in growth media) were administered and tracked simultaneously upon pipetting of 50 µL tracer solution into 50 µL bare culture media in the media layer (2x final conc.). A 1×5 tilescan (350 µm x 1750 µm – center tile containing the 300 µm barrier) was taken at substrate-height every 10 minutes for 12 hrs with T=0 min directly after application. Diffusion was quantified at selected timepoints by generating intensity profiles for rectangular ROIs right outside the microtunnels (somatic and axonal compartments) and in the bulk (somatic). Time-dependent tracer loss to the somatic channel was then accounted for by measuring the fluorescent intensity development in both compartments and normalized as a % of the highest value measured axonally (at T=12 hrs).

### Calcium imaging

Neuron-cancer cocultures were transduced as described in “AAV1-eGFP and AAV1-GCaMP6f transduction”. Adeno-associated virus expressing GCaMP6f as Ca2+ indicator was imaged before and immediately after application of inflammatory/excitatory compounds using a spinning disk Zeiss Cell Observer® System (20x/63x objective) and processed in FIJI. At 5-6 days in culture, cocultures were placed inside the 384 well microplate adaptor and left to acclimate. ROI (350 µm x 350 µm) was selected by qualifying the presence of innervated cancer spheroids at various heights within the gel layer before stimulation with either 4 µM Bradykinin (Tocris/Biotechne, #3004) or 1 µM Capsaicin (Tocris/Biotechne, #0462). Acquisition time was set at ∼2 frames per second.

### MEA-based electrophysiological recordings

Electrophysiology of 3D innervating fibers was recorded using a USB-MEA 120-System (Multi Channel Systems MCS GmbH, Germany) at 4 – 6 days in culture for DRG monocultures and at 6 days in culture for cancer cocultures. During experiments, a customized MEA incubation chamber (Okolab Srl, Italy) was used to continuously monitor temperature, CO2 and humidity. The temperature was set to 36◦C, humidity at 85% and CO2 at 5%. MEAs were allowed to acclimate inside the MEA chamber before applying agonists and inflammatory mediators to the axonal compartment. Agonists were applied 30 seconds into a recording to identify spontaneously active electrodes. Inflammatory mediators were applied 5 minutes before recording. Recordings were acquired using MC_Rack v. 4.6.2 (Multichannel Systems MCS GmbH, Germany) and filtered using a fourth-order bandpass filter (60-6000 Hz), before processing with NeuroExplorer (version 5.300) for spike detection and further analysis.

### Statistical analysis

Statistical analysis was performed on the collected data using GraphPad Prism 8 developed by GraphPad Software Inc. (Boston, USA). Degree of normality of the data distribution was assessed both visually and quantitatively by employing quantile-quantile plots, frequency histograms, and Shapiro-Wilks tests. Within the neurite outgrowth assay domain, conditions are represented as bar graphs with mean ± SEM. Differences were assessed by Brown-Forsythe and Welch’s ANOVA test with post-hoc test Dunnett T3 (n<50) for comparison of means. Electrophysiological data were represented as either bar graphs comparing conditions or connected-points for temporal dynamics with mean ± SEM. Differences were assessed similarly by either Brown-Forsythe and Welch’s ANOVA with post hoc test Dunnett T3 (n< 50) or Welch’s unpaired t-tests for comparison of means. Asterisks denote statistical significance as follows: ∗p < 0.05; ∗∗p < 0.01; ∗∗∗p < 0.001; ∗∗∗∗p < 0.0001.

## Acknowledgements

We thank Rosanna Toscano (FEMTOprint, Muzzano, CH) for fruitful discussions on glass microfluidics, Anna-Lena Keller (Tumor Biology, NMI) for supplying the human cancer cell lines and Michael Mierzejewski, Peter Jones (Bio-medical Micro and Nano Engineering, NMI) and Eduardo Brás (Organ-on-Chip, NMI) for technical assistance. We also thank Frank Machnow (NMI) for comprehensive 3D modelling and Martin Kriebel (Molecular Biology & Neurobiology, NMI) for confocal microscopy assistance.

## Funding

PC acknowledges funding from the State Ministry of Baden-Württemberg for Economic Affairs, Labour and Housing Construction. MvdM and PC acknowledge funding from the European Union Horizon 2020 Framework Programme for Research an Innovation under the Marie Sklodowska-Curie Grant Agreement No. 814244 (BONEPAIN II). PC acknowledges funding from Eureka’s Eurostars Programme under grant agreement 115217 (NEUROCHIP). FE acknowledges funding from the Ministry of National Education of Turkey.

## Author Contributions

M.M. and P.C. designed research; M.M. and P.C. performed biological and microfabrication research; F.E. aided microfabrication. A. L. and S. M. created glass devices, M.M. and P.C. analyzed data; M.M., P.L. and P.C. wrote the paper.

## Competing Interest Statement

P.C. is named as inventor on patent application WO2019115320A1 (“Device for the examination of neurons”) filed by NMI Natural and Medical Sciences Institute at the University of Tübingen.

## Notes

### Competing Interest Statement

Paolo Cesare is named as inventor on patent applications EP3494877 and WO2019115320 (Device for the examination of neurons) filed by NMI Natural and Medical Sciences Institute at the University of Tuebingen.

## References

[1] D. Park, J. Lee, J. J. Chung, Y. Jung, and S. H. Kim, "Integrating Organs-on-Chips: Multiplexing, Scaling, Vascularization, and Innervation," Trends Biotechnol, vol. 38, no. 1, pp. 99–112, Jan 2020, doi: 10.1016/j.tibtech.2019.06.006.

[2] P. S. Rajendran et al., "Identification of peripheral neural circuits that regulate heart rate using optogenetic and viral vector strategies," Nat Commun, vol. 10, no. 1, p. 1944, Apr 26 2019, doi: 10.1038/s41467-019-09770-1.

[3] S. Takahashi et al., "Homeostatic pruning and activity of epidermal nerves are dysregulated in barrier-impaired skin during chronic itch development," Sci Rep, vol. 9, no. 1, p. 8625, Jun 13 2019, doi: 10.1038/s41598-019-44866-0.

[4] X. Guo et al., "Tissue engineering the mechanosensory circuit of the stretch reflex arc with human stem cells: Sensory neuron innervation of intrafusal muscle fibers," Biomaterials, vol. 122, pp. 179–187, Apr 2017, doi: 10.1016/j.biomaterials.2017.01.005.

[5] N. Vysokov, S. B. McMahon, and R. Raouf, "The role of Na(V) channels in synaptic transmission after axotomy in a microfluidic culture platform," Sci Rep, vol. 9, no. 1, p. 12915, Sep 9 2019, doi: 10.1038/s41598-019-49214-w.

[6] R. Atmaramani et al., "Investigating the Function of Adult DRG Neuron Axons Using an In Vitro Microfluidic Culture System," Micromachines (Basel), vol. 12, no. 11, Oct 27 2021, doi: 10.3390/mi12111317.

[7] E. Neto et al., "Osteoclast-derived extracellular vesicles are implicated in sensory neurons sprouting through the activation of epidermal growth factor signaling," Cell Biosci, vol. 12, no. 1, p. 127, Aug 14 2022, doi: 10.1186/s13578-022-00864-w.

[8] H. A. Enright et al., "Long-term non-invasive interrogation of human dorsal root ganglion neuronal cultures on an integrated microfluidic multielectrode array platform," Analyst, vol. 141, no. 18, pp. 5346–57, Sep 21 2016, doi: 10.1039/c5an01728a.

[9] A. Klusch, C. Gorzelanny, P. W. Reeh, M. Schmelz, M. Petersen, and S. K. Sauer, "Local NGF and GDNF levels modulate morphology and function of porcine DRG neurites, In Vitro," PLoS One, vol. 13, no. 9, p. e0203215, 2018, doi: 10.1371/journal.pone.0203215.

[10] A. J. Clark et al., "Functional imaging in microfluidic chambers reveals sensory neuron sensitivity is differentially regulated between neuronal regions," Pain, vol. 159, no. 7, pp. 1413–1425, Jul 2018, doi: 10.1097/j.pain.0000000000001145.

[11] B. Molina-Martinez et al., "A multimodal 3D neuro-microphysiological system with neurite-trapping microelectrodes," Biofabrication, vol. 14, no. 2, Jan 24 2022, doi: 10.1088/1758-5090/ac463b.

[12] P. W. Mantyh, "Mechanisms that drive bone pain across the lifespan," Br J Clin Pharmacol, vol. 85, no. 6, pp. 1103–1113, Jun 2019, doi: 10.1111/bcp.13801.

[13] J. M. Jimenez-Andrade et al., "Pathological sprouting of adult nociceptors in chronic prostate cancer-induced bone pain," J Neurosci, vol. 30, no. 44, pp. 14649–56, Nov 3 2010, doi: 10.1523/JNEUROSCI.3300-10.2010.

[14] S. M. Gysler and R. Drapkin, "Tumor innervation: peripheral nerves take control of the tumor microenvironment," J Clin Invest, vol. 131, no. 11, Jun 1 2021, doi: 10.1172/JCI147276.

[15] Q. Zheng, D. Fang, J. Cai, Y. Wan, J. S. Han, and G. G. Xing, "Enhanced excitability of small dorsal root ganglion neurons in rats with bone cancer pain," Mol Pain, vol. 8, p. 24, Apr 3 2012, doi: 10.1186/1744-8069-8-24.

[16] A. P. Bloom et al., "Breast cancer-induced bone remodeling, skeletal pain, and sprouting of sensory nerve fibers," J Pain, vol. 12, no. 6, pp. 698–711, Jun 2011, doi: 10.1016/j.jpain.2010.12.016.

[17] R. Haroun, J. N. Wood, and S. Sikandar, "Mechanisms of cancer pain," Front Pain Res (Lausanne), vol. 3, p. 1030899, 2022, doi: 10.3389/fpain.2022.1030899.

[18] Y. F. Zhu et al., "Differences in electrophysiological properties of functionally identified nociceptive sensory neurons in an animal model of cancer-induced bone pain," Mol Pain, vol. 12, 2016, doi: 10.1177/1744806916628778.

[19] E. J. Huang and L. F. Reichardt, "Neurotrophins: roles in neuronal development and function," Annu Rev Neurosci, vol. 24, pp. 677–736, 2001, doi: 10.1146/annurev.neuro.24.1.677.

[20] T. J. Price et al., "Treatment of trigeminal ganglion neurons in vitro with NGF, GDNF or BDNF: effects on neuronal survival, neurochemical properties and TRPV1-mediated neuropeptide secretion," BMC Neurosci, vol. 6, p. 4, Jan 24 2005, doi: 10.1186/1471-2202-6-4.

[21] E. Cao, M. Liao, Y. Cheng, and D. Julius, "TRPV1 structures in distinct conformations reveal activation mechanisms," Nature, vol. 504, no. 7478, pp. 113–118, 2013, doi: 10.1038/nature12823.

[22] D. Albo et al., "Neurogenesis in colorectal cancer is a marker of aggressive tumor behavior and poor outcomes," Cancer, vol. 117, no. 21, pp. 4834–45, Nov 1 2011, doi: 10.1002/cncr.26117.

[23] G. Gasparini et al., "Nerves and Pancreatic Cancer: New Insights into a Dangerous Relationship," Cancers (Basel), vol. 11, no. 7, Jun 26 2019, doi: 10.3390/cancers11070893.

[24] A. Horn, Dahl, O., & Morild, I., "Venous and neural invasion as predictors of recurrence in rectal adenocarcinoma," Diseases of the colon & rectum, 1991.

[25] R. E. Stopczynski et al., "Neuroplastic changes occur early in the development of pancreatic ductal adenocarcinoma," Cancer Res, vol. 74, no. 6, pp. 1718–27, Mar 15 2014, doi: 10.1158/0008-5472.CAN-13-2050.

[26] D. Huh, B. D. Matthews, A. Mammoto, M. Montoya-Zavala, H. Y. Hsin, and D. E. Ingber, "Reconstituting organ-level lung functions on a chip," Science, vol. 328, no. 5986, pp. 1662–8, Jun 25 2010, doi: 10.1126/science.1188302.

[27] S. Das et al., "Innervation: the missing link for biofabricated tissues and organs," NPJ Regen Med, vol. 5, p. 11, 2020, doi: 10.1038/s41536-020-0096-1.

[28] R. B. Campenot, "Local control of neurite development by nerve growth factor," PNAS, 1977.

[29] D. Am., "The emerging generality of neurotrophic hypothesis," Trends in Neurosciences, 1988.

[30] X.-Q. L. Shu, A.; Mendell, L. M.*, "Effects of trkB and trkC neurotrophin receptor agonists on thermal nociception a behavioral and electrophysiological study," PAIN, 1999.

[31] R. A. M. IS. D. Davis, and J. N. Campbell, "Chemosensitivity and Sensitization of Nociceptive Afferents that Innervate the Hairy Skin of Monkey," Journal of Neurophysiology, 1993.

[32] X. Li et al., "Targeting tumor innervation: premises, promises, and challenges," Cell Death Discov, vol. 8, no. 1, p. 131, Mar 25 2022, doi: 10.1038/s41420-022-00930-9.

[33] X. Chu et al., "Blocking Cancer-Nerve Crosstalk for Treatment of Metastatic Bone Cancer Pain," Adv Mater, vol. 34, no. 17, p. e2108653, Apr 2022, doi: 10.1002/adma.202108653.

[34] D. A. Silverman, V. K. Martinez, P. M. Dougherty, J. N. Myers, G. A. Calin, and M. Amit, "Cancer-Associated Neurogenesis and Nerve-Cancer Cross-talk," Cancer Res, vol. 81, no. 6, pp. 1431–1440, Mar 15 2021, doi: 10.1158/0008-5472.CAN-20-2793.

[35] Z. Gil et al., "Paracrine regulation of pancreatic cancer cell invasion by peripheral nerves," J Natl Cancer Inst, vol. 102, no. 2, pp. 107–18, Jan 20 2010, doi: 10.1093/jnci/djp456.

[36] I. E. Demir, H. Friess, and G. O. Ceyhan, "Neural plasticity in pancreatitis and pancreatic cancer," Nat Rev Gastroenterol Hepatol, vol. 12, no. 11, pp. 649–59, Nov 2015, doi: 10.1038/nrgastro.2015.166.

[37] L. S. Caroline Brunetto de Fariasa, Rodrigo Cruz Limaa, Flávio Kapczinskic, Gilberto Schwartsmann and Rafael Roesler, "Reduced NGF Secretion by HT-29 Human Colon Cancer Cells Treated with a GRPR Antagonist," Protein & Peptide Letters, 2009.

[38] H. F. Zhao-wen Zhu, Li Wang, Thomas Bogardus, Murray Korc, Jorg Kleeff, and Markus W. Buchler, "Nerve Growth Factor Exerts Differential Effects on the Growth of Human Pancreatic Cancer Cells," Clinical Cancer Research, 2001.

[39] H. Wang et al., "Role of the nervous system in cancers: a review," Cell Death Discov, vol. 7, no. 1, p. 76, Apr 12 2021, doi: 10.1038/s41420-021-00450-y.

[40] Y. Bellouard, "The Femtoprint Project," Journal of Laser Micro/Nanoengineering, vol. 7, no. 1, pp. 1–10, 2012, doi: 10.2961/jlmn.2012.01.0001.

[41] J. N. Sleigh, G. A. Weir, and G. Schiavo, "A simple, step-by-step dissection protocol for the rapid isolation of mouse dorsal root ganglia," BMC Res Notes, vol. 9, p. 82, Feb 11 2016, doi: 10.1186/s13104-016-1915-8.

