## Supplementary Figures for "Cancer-mediated Axonal Guidance of Sensory Neurons in a Microelectrode-based Innervation MPS"

### Supplemental Information

### Supplementary Figures

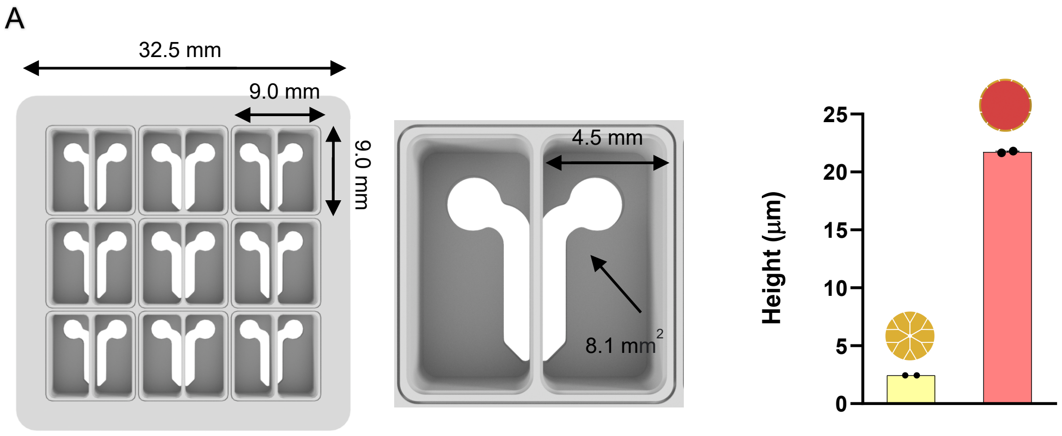


**Fig. S1 INV-MPS overview.** (A) Important parameters given as input for SLE of fused silica microfluidic chips (B) Stylus profilometry of subsequent photoresist layers constituting a CME. Data in B is represented as mean ± SEM.

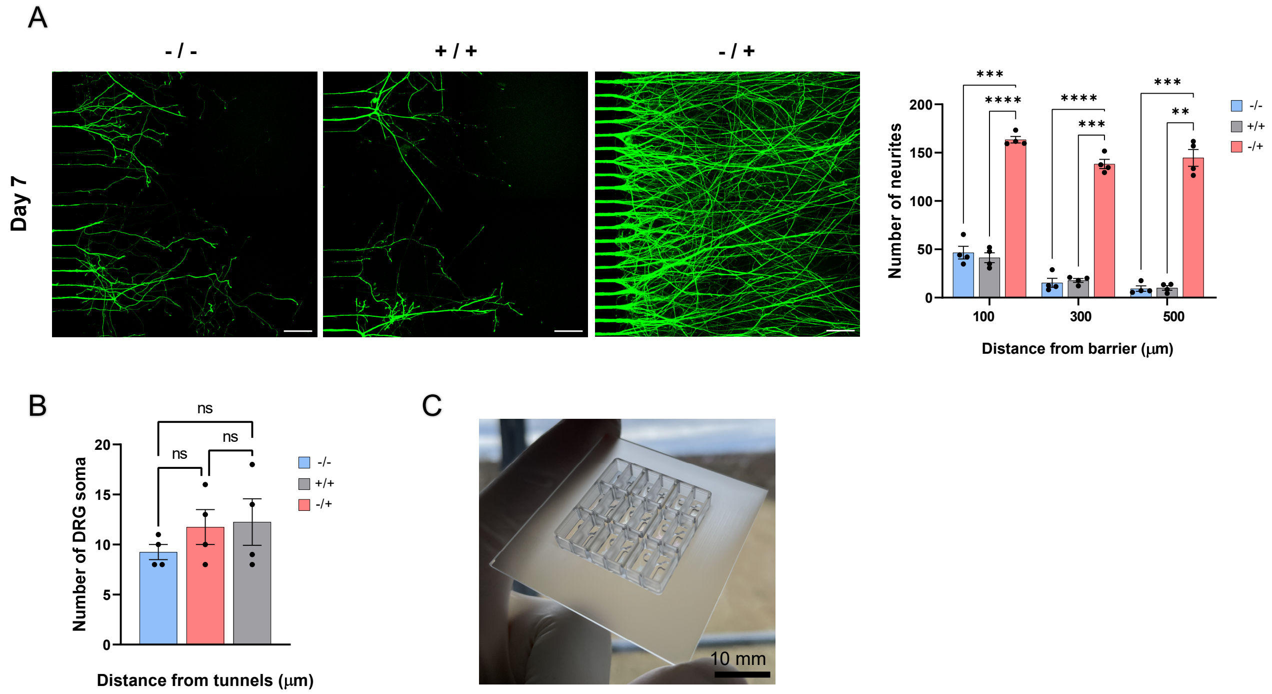


**Fig. S2 Axonal outgrowth and survival.** (A) DRG neurons extending in 3D inside the axonal compartment after 7 days in culture with quantification. (n = 4 wells) (Scale bar: 75 µm.) (B) Comparison of neuronal survival inside the somatic compartment by counting eGFP-fluorescent DGR soma. (n = 4 wells) (C) SLE INV-MPS without electrodes used for growth quantification and diffusion. In A and B, bar graphs represent mean ± SEM and significant differences between mean values of groups are displayed as asterisks: ∗P < 0.05; ∗∗p < 0.01; ∗∗∗p < 0.001; ∗∗∗∗p < 0.0001; Brown-Forsythe and Welch's ANOVA test.

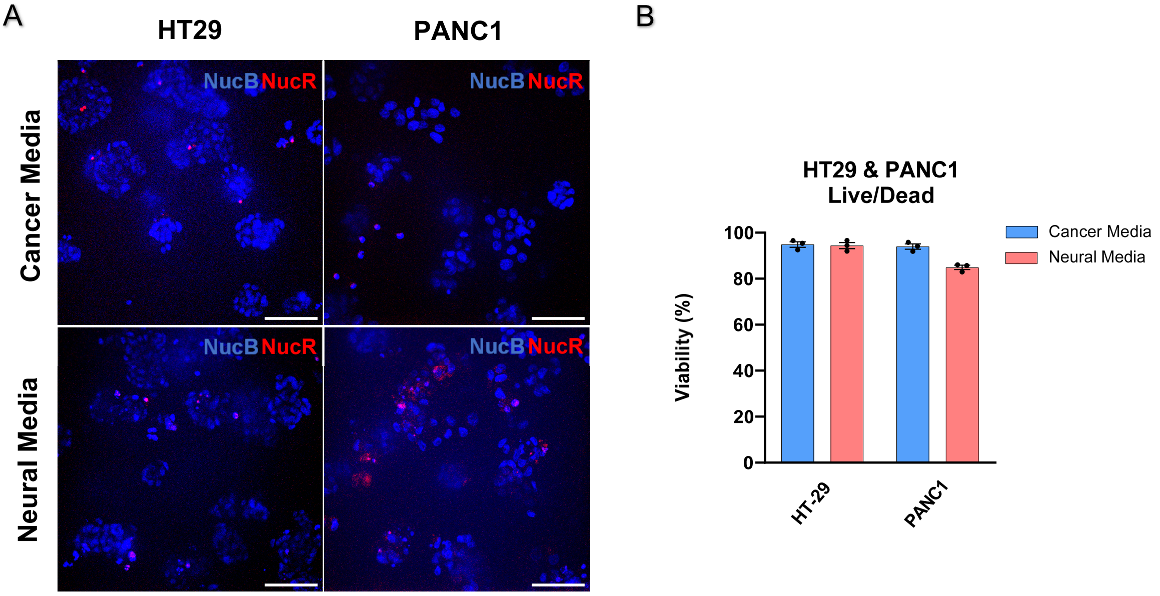


**Fig. S3 Cancer viability study.** (A) Cancer spheroids are imaged after 6 days in culture using dyes staining nuclei (NucB) and dead cells (NucR). HT29 and PANC1 are either cultivated in their own media or neuronal media (Scale bar: 75 µm.). (B) Live/dead quantification (Total Cells - Dead cells/Total Cells * 100%). Bar graphs represent mean ± SEM (n = 3 wells).

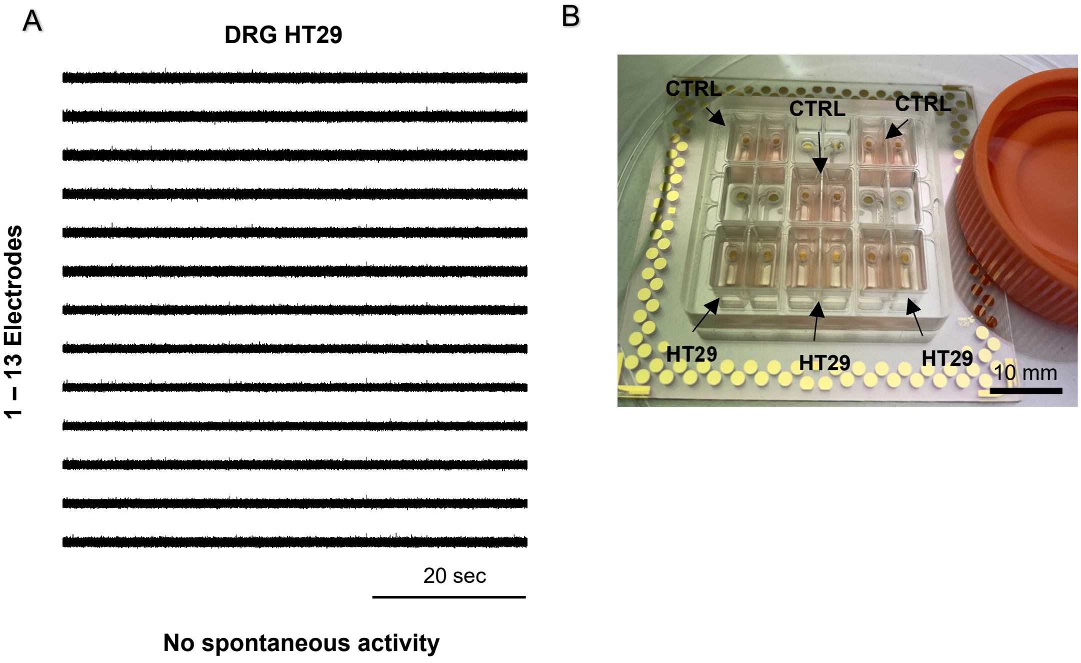


**Fig. S4 Sensory innervation of cancer spheroids case study.** (A) Spheroids do not induce spontaneous activity in DRG neurons that extend into the axonal compartment (B) Overview of INV-MPS setup on a recording day

### Supplementary Videos

**Movie. S1. 3D-surface rendering of DRG terminals.** Nerve fibers project into 3D space after barrier exit. Growing somatic/visceral tissues in between these protruding fibers could replicate innervation. Dimensions 600 × 450 × 300 µm. (Scale bar: 100 µm).

**Movie. S2. Z-stack of GFP-transduced DRG inside the somatic compartment.** Soma can be seen homogeneously distributed across the somatic compartment in 3D space with CMEs observable at the bottom. CMEs are 250.0 µm wide and interspaced 600.0 µm center-to-center. Dimensions 700 × 700 × 300 µm. (Scale bar: 100 µm.)

**Movie. S3. 3D rendering of GFP-transduced DRG inside an INV-MPS.** Soma and neurites are spatially separated by the micro tunneled barrier. Neurites are seen extending into 3D space after barrier exit. Dimensions 1450 × 750 × 300 µm. (Scale bar: 150 µm.)

**Movie. S4. Z-stack of GFP-transduced DRG** (Green) **innervating NucBlue** (Hoechst) **stained HT29 cancer spheroids** (Blue). Stack was used to 3D-surface render Movie. S6. Dimensions 400 x 400 x 200 µm. (Scale bar: 25 µm.)

**Movie. S5. Z-stack of GFP-transduced DRG** (Green) **innervating NucBlue** (Hoechst) **stained PANC1 cancer spheroids** (Blue). Stack was used to 3D-surface render Movie. S7. Dimensions 400 x 400 x 200 µm. (Scale bar: 25 µm.)

**Movie. S6. 3D-surface renderings of DRG sensory neurites** (Green**) innervating HT29 cancer spheroids** (Blue). Dimensions 400 x 400 x 200 µm. (Scale bar: 70 µm.)

**Movie. S7. 3D-surface renderings of DRG sensory neurites** (Green) **innervating PANC1 cancer spheroids** (Blue). Dimensions 400 x 400 x 200 µm. (Scale bar: 50 µm.)

**Movie. S8. Sensory innervation of cancer spheroids inside the INV-MPS driving excitability.** Cancer spheroids (Blue) stimulate the depolarization of nerve terminals (Green). Action potentials generated upon compound application travel through the barrier and are transmitted into CMEs.

**Movie. S9. Timeseries of GCaMPf6-transduced DRG neurites showing $Ca^{2+}$ transients while enveloping a HT29 cancer spheroid.** Real-time experiment ran at 2.1 frames per second. Application; Bradykinin 4.0 µM. (Scale bar: 25 µm.)

**Movie. S10. Timeseries of GCaMPf6-transduced DRG neurites showing $Ca^{2+}$ transients while enveloping a HT29 cancer spheroid.** Real-time experiment ran at 1.5 frames per second. Application; Capsaicin 1.0 µM. (Scale bar: 25 µm.)
